# Açai (*Euterpe oleracea Mart*) modulates oxidative stress and inflammation by NF-κB inactivation and Nrf2 up-regulation in experimental diabetes

**DOI:** 10.1101/2022.02.14.480447

**Authors:** Deyse Yorgos de Lima, Adelson Marçal Rodrigues, Margaret Gori Mouro, Elias Jorge Muniz Seif, Giovana Rita Punaro, Elisa Mieko Suemitsu Higa

## Abstract

To evaluate the effects of açai extract (EA) on oxidative stress and inflammation induced by high glucose in cultured mouse immortalized mesangial cells (MiMC) and diabetic rats. MiMC cell viability and proliferation were determined by MTT. Extracellular and intracellular nitric oxide (NO) and intracellular ROS were also measured. The cell proteins were extracted for analysis of catalase, Nrf2, p-Nrf2, SOD-1, SOD-2, iNOS, NF-κB, p-NF-κB and TNF-α expression. Male, adult Wistar rats were distributed into 3 groups: control (CTL) and diabetic (DM) rats who received water and DMEA and received 1 mL/day EA (200 mg/kg) via gavage for 8 consecutive weeks. After treatment with EA, metabolic profile, renal function and thiobarbituric acid reactives substances (TBARS) levels were evaluated, and kidneys were collected for qualitative histological analysis. EA maintained cell viability above 90% in all groups; it decreased proliferation in the HG group, both significant. NO levels, ROS generation, iNOS, NF-κB, p-NF-κB and TNF-α expression were reduced significantly after 72 h of EA treatment, with significant increases for all antioxidants studied. DMEA *vs* DM showed a significant increase in body weight, improved kidney function and reduced TBARS excretion. EA treatment decreased proliferation, oxidative stress and inflammation in MiMC, and although açai did not decrease fasting glucose, it recovered the body weight and delayed the decline of renal function in the diabetic animals, suppressing the signaling of inflammatory mediators via NF-κB inactivation and increasing all antioxidants studied by upregulating the Nrf2 response pathway.

## 1. Introduction

Diabetes mellitus (DM) is a chronic nontransmissible and inflammatory disease characterized by hyperglycemia, which leads, among other complications, to microvascular damage such as neuropathy, retinopathy and diabetic nephropathy (DN) [1]. DN is represented by expansion of mesangial extracellular matrix (ECM), resulting in gradual reduction of glomerular filtration [2]. Mesangial cells are responsible for the regulation of glomerular filtration, phagocytosis of immune complexes and ECM production; when stimulated by immunological or inflammatory factors, they can generate cytokines, chemokines and NO [3]; the latter is synthesized by the enzyme nitric oxide synthase (NOS), and its main action is the relaxation of blood vessels [4]. It is known that glomerular mesangial cell injury plays a crucial role in the development of DN, since these cells are more susceptible to apoptosis and injury induced by hyperglycemia [5].

Hyperglycemia leads to the synthesis of ROS, which act in the degradation of macromolecules [6]. This occurs by means of a nonenzymatic and irreversible reaction, forming advanced glycation end products (AGEs) [7]; these, in turn, activate the nuclear transcription factor NF-κB [8, 9] that stimulates proinflammatory cytokines and tumoral necrosis factor-α (TNF-α) [10].

Oxidative stress is a critical factor in the pathophysiology of DN and is one of the main mechanisms of this disease [11]. A previous study in our Laboratory showed that antioxidants such as cupuaçu had a beneficial effect on diabetic models [12]. Thus, the use of antioxidants as a therapeutic strategy could alleviate cellular damage in DM.

Experimental models of diabetes induced in rats have been widely used by researchers worldwide because animal models have clinical, laboratory and histopathological similarities with human diabetes [13]. Several studies have shown that changes in diet and rich foods in antioxidants, tested in animal models, can improve kidney function and clinical markers related to diabetes and inflammation [14-17].

Açai is a natural tropical fruit from the Amazon that originates from the palm açai tree (Euterpe oleracea Mart) and is consumed in different parts of the world [18]. It is considered a high-caloric food due to its high lipid content and high nutritive content because it is rich in protein and α-tocopherol (vitamin E), fibers, manganese, copper, boron, calcium, magnesium, potassium and chromium [19, 20].

Previous studies have shown that the use of antioxidants, such as kefir and N-acetylcysteine, contributed to reducing hyperglycemia and ROS, improving renal function in diabetic animals, and delaying the effects of DN [15, 21]. The aim of this study was to evaluate the effects of EA on oxidative stress and inflammation induced by high glucose in MiMC and diabetic rats.

## 2. Material and methods

### 2.1. Assays *in vitro*

#### 2.1.1. Açai extract (EA)

Freeze-dried açai (100 mg) from Liotecnica Alimentos LTDA (Sao Paulo, Brazil, Cod. 120.044.032) was diluted in 1 mL of phosphate buffered saline (PBS, 0.01 M, pH 7.4) to a concentration of 100 mg/mL, mixed and centrifuged at 2000 rpm [22]. The extract was filtered through a 0.22 μm syringe. To attain the concentrations used for treatment, it was further diluted in PBS to 500, 100 and 50 µg/mL, and the product was kept at -80°C [23, 24]. Each 10 g of dry weight contained 5.4 g of lipids (3.0 g of omega-9 and 0.5 g omega-6), 1.0 g of protein, 0.5 g of carbohydrates, 2.7 g of fibers, and 54.1 kcal.

##### 2.1.1.1. Total polyphenol content

The total flavonoid content (TFC) was estimated by the aluminum chloride colorimetric method; for quantitative analysis, quercetin (QUE) (concentration range of 10–100 μg/mL) was chosen as the standard compound, and the measurements were performed at 435 nm. TFC was measured in QUE/g/100 g dry weight of açai (R^2^ = 0.98) [25].

##### 2.1.1.2. Total flavonoid content

Total polyphenol content was determined by the Folin-Ciocalteu method [26]. Different concentrations of gallic acid (Sigma-Aldrich, Saint Louis, MO, USA) were used to construct a standard curve for quantifying total polyphenols, and the values were expressed in mg gallic acid equivalents (GAE) (0-1,000 mg/L) per 100 g of lyophilized açai, (R^2^ = 0.97) [27].

##### 2.1.1.3. Antioxidant activity

The antioxidant activity of freeze-dried açai was determined by the 2,2-diphenyl-1-picrylhydrazyl assay (DPPH), which evaluates the ability of a substance to scavenge the free radical DPPH; we used the IC_50_ parameter, which represents the concentration of the material in question necessary to inhibit 50% of free radicals, measured at 515 nm and converted into the percentage of antioxidant activity, using the following formula: AA% = 100 – {[(Abs _sample_ – Abs _blank_) × 100]/Abs _control_}

The EC_50_ of gallic acid standard antioxidant was 0.269 µg/mL. All analyses were carried out in triplicate [28].

#### 2.1.2. Mesangial cell culture

MiMC SV40 MES 13 (CRL 1927 - ATCC) was provided by the Nephrology Division – UNIFESP/EPM. The cells were grown and kept in a 95% air and 5% CO2 humidified environment at 37 °C in Dulbecco’s modified Eagle’s medium (DMEM) and F12 (3:1) containing 5% fetal bovine serum (FBS) and penicillin (50 U/mL)/streptomycin (50 mg/mL). All products were obtained from Gibco-Life Technology (Sao Paulo, Brazil). The medium was replaced every 48 h. All experiments were performed with cells between the 10^th^ and 20^th^ passages. The ideal time (72 h) for treatment was determined according to the time response curve of proliferation and viability of MiMCs exposed to high glucose and/or açai. Recent studies have used a dose of 30 mM as a high glucose concentration to mimic diabetes [29, 30].

At semiconfluence of approximately 60-70% (ideal cell density to avoid overgrowth, as previously tested, in our Laboratory), the cells were cultured in FBS-free medium for 24 h, and after this period, the medium was replaced, and the cells were treated for 24, 48 or 72 h with 0.5% FBS according to the experimental groups: normal glucose group (NG), which was cultured in medium containing a standard concentration of 6.7 mM D-glucose; high glucose group (HG), cultured with medium containing D-glucose at a final concentration of 30 mM (Sigma, Sao Paulo, Brazil); and the osmolarity control group, which was cultured in medium supplemented with mannitol (MA, Sigma, Sao Paulo, Brazil) at a final concentration of 30 mM.

#### 2.1.3. Cell viability and proliferation

The MiMCs were cultured with 5% FBS in 96-well culture plates at a concentration of 5 × 104 cells/mL per well. At semiconfluence (60-70%), the cells were exposed to FBS-free medium for 24 h for synchronization. Then, the medium was replaced with DMEM containing 0.5% FBS in conditions of NG, MA, HG or HG plus 500, 100 or 50 µg/mL açai extract for 72 h. After this time, the cells were trypsinized and centrifuged, the supernatant was discarded, and the cells were resuspended in 1 mL of PBS. Proliferation was evaluated by methyl thiazole tetrazolium assay (MTT, Sigma, Sao Paulo, Brazil). Optical density (OD) was measured with a microplate reader at a wavelength of 570 nm [31]. The OD of the NG cells was assigned by a relative value of 100. The results were obtained from averages of three independent experiments performed in octuplicate wells.

#### 2.1.4. NO measurement

Twelve-well culture plates with a concentration of 7 × 10^3^ cells/mL per well of MiMC were treated according to their respective groups. After the treatment, the supernatant was collected and stored in a freezer at -20 °C. The NO levels in the supernatant were measured by chemiluminescence using a Nitric Oxide Analyzer (NOA 280, Sievers Instruments Inc, USA), a high-sensitivity detector for measuring NO (∼1 pmol), which is based on the gas-phase chemiluminescent reaction between NO and ozone. The results were obtained from three independent experiments performed in triplicate wells [32].

#### 2.1.5. Fluorescence of intracellular NO

A concentration of 6 × 10^3^ cells was loaded with 5 mM 4-amino-5-methylamino-2’,7’- difluorofluorescein diacetate (DAF-FM, Molecular Probes, Thermo Fisher, Sao Paulo, Brazil) after 72 h of treatment. DAF-FM is a reagent that is used to detect and quantify low concentrations of NO. Ratios of green (NO-DAF-FM) to blue with 4,6-diamidino-2-fenilindol (DAPI) fluorescence were tabulated from three images of fields containing 15 or more cells, and the ratios were normalized by the control for each experiment [33]. The intensity of the fluorescence was quantified by ImageJ software (US National Institutes of Health, MD, USA).

#### 2.1.6. Assessment of intracellular ROS

Intracellular ROS levels were measured using the nonfluorescent probe 2,7-dichlorodihydrofluorescein diacetate (DCFH-DA, Sigma, Sao Paulo, Brazil), which is converted into 2’,7’ dichlorodihydrofluorescein fluorescent (DCF) in the presence of intracellular hydrogen peroxide and peroxidases. The MiMCs were incubated in DCFH-DA (9 μM) at 37 °C for 30 min, the medium was then removed, and the cells were washed with PBS. The fluorescence was measured using a fluorescence plate reader (Synergy HT, Biotek, USA); excitation was read at 480 nm, and emission was detected at 520 nm [34]. Relative ROS production was expressed as the mean fluorescence intensity of DCF. The results were obtained from three independent assays performed in octuplicate wells.

#### 2.1.7. Protein content

The cells (1 × 10^5^ cells/mL) were cultured in 100 mm Petri dishes and treated according to the groups cited above for 72 h. After this time, the cells were lysed with RIPA buffer and protease inhibitor. The protein was concentrated with ultra-filter 0.5 (Millipore, Sao Paulo, Brazil), and it was determined by BCA protein assay (Sigma, Sao Paulo, Brazil). A total of 10 µg protein concentrate was used in a 10% or 12% polyacrylamide gel and transferred to a nitrocellulose membrane. Nonspecific binding was blocked with 10% nonfat dry milk in a pH 7.5 TBS-T buffer followed by washing in the same buffer at room temperature. The membranes were then incubated overnight at 4 °C with primary antibodies against catalase (Sigma, Sao Paulo, Brazil), Nrf2, p-Nrf2 (Ser40), SOD-1, SOD-2, iNOS, NF-κB p65, p-NF-κB p65, TNF-α and housekeeping actin (Santa Cruz Biotechnology Inc., CA, USA). The specific protein bands were visualized with immobilon western chemiluminescent HRP substrate (Millipore, Sao Paulo, Brazil), captured by Alliance 4.7 (Uvitec, Cambridge, UK) and quantified by ImageJ software for analysis of Western blots. The results were obtained from three independent assays performed in duplicate [35].

### 2.2. Assays in vivo

#### 2.2.1. Animals

For in vivo experiments, freeze-dried açai (Liotecnica Alimentos LTDA) (Sao Paulo, Brazil, code 120.044.032) was weighed three times a week according to the animals’ body weight, protected from light and kept at 4 °C until use. At the time of gavage, the açai was diluted in 1 mL of filtered water, homogenized and given to the animals. All these procedures were performed in a dark environment to prevent oxidation of the compounds. The animal protocol was approved by the Ethics Committee in Research at Federal University of Sao Paulo, under number #6729100418.

Male Wistar rats (Rattus norvegicus), 7 weeks old, weighing 170-210 g were obtained from the Center for Experimental Model Development (CEDEME) of the Universidade Federal of Sao Paulo - UNIFESP. All procedures were carried out in accordance with the ARRIVE guidelines - Animal Research: Reporting of In Vivo Experiments [36]; the guides for Anesthesia, Analgesia, Euthanasia and model for monitoring laboratory animals - CEUA / UNIFESP (Version 2019), and the Euthanasia Practice Guidelines of the (National Council for Animal Experimentation Control) CONCEA - Normative Resolution n° 37. They were maintained with a controlled temperature of 22 ± 2 °C and environment with a regular period of light and dark 12:12 h, receiving standard chow and water ad libitum. The rats were distributed into three groups: CTL (control); DM (diabetic) and DMEA (diabetic treated with açai), with n = 4, CTL group, and 5, for DM group.

#### 2.2.2. Nephrectomy

To accelerate the development of DN, unilateral nephrectomy was performed in all animals. After trichotomy and skin cleansing with 70% ethyl alcohol and 50% polyvinylpyrrolidone solution, the animals were anesthetized with ketamine hydrochloride (65 mg/kg, Sesp, Sao Paulo, Brazil) and xylazine hydrochloride (9 mg/kg, Rhobifarma, Sao Paulo, Brazil), both i.v. Then, a small incision (± 2 cm) was made in the lumbar region, and the left kidney was removed. Nonsteroidal anti-inflammatory meloxicam (2 mg/kg, Maxicam, Sao Paulo, Brazil) was administered shortly after the procedure for 3 days, once a day, subcutaneously.

#### 2.2.3. DM induction

After a few days of adaptation, 8-week-old rats received a single intravenous dose of 60 mg/kg streptozotocin (STZ, Sigma Chemical, MO, USA) diluted in STZ-vehicle (cold citrate buffer, 0.1 M, pH 4.5). Control rats received STZ-vehicle. After 48 h of induction, glycemia was checked through a glucometer in a blood sample, which was collected from the tail vein. Diabetes was determined when fasting glycemia was 200 mg/dL. The DMEA received extract of açai (EA) beginning on the 4th day after induction of diabetes. EA (1 mL/day) was given for 8 consecutive weeks via gavage at a concentration of 200 mg/kg/day. The CTL and DM groups received EA-vehicle (water).

#### 2.2.4. Metabolic profile and euthanasia

The animals were individually placed in metabolic cages (Tecniplast, Buguggiate, Italy) for 24 h, receiving standard chow and water ad libitum before and at the end of the 8th week of the protocol. Three hours of fasting glucose measurement, diuresis (mL/24 h), water (mL/24 h) and chow (mg/24 h) were obtained. The animals were euthanized with 90 mg/kg ketamine chloridrate and 18 mg/kg xylazine chloridrate intraperitoneally (according to the guides already mentioned), followed by an incision of the diaphragm, and the right kidney was removed. There were no adverse events in the experimental protocol.

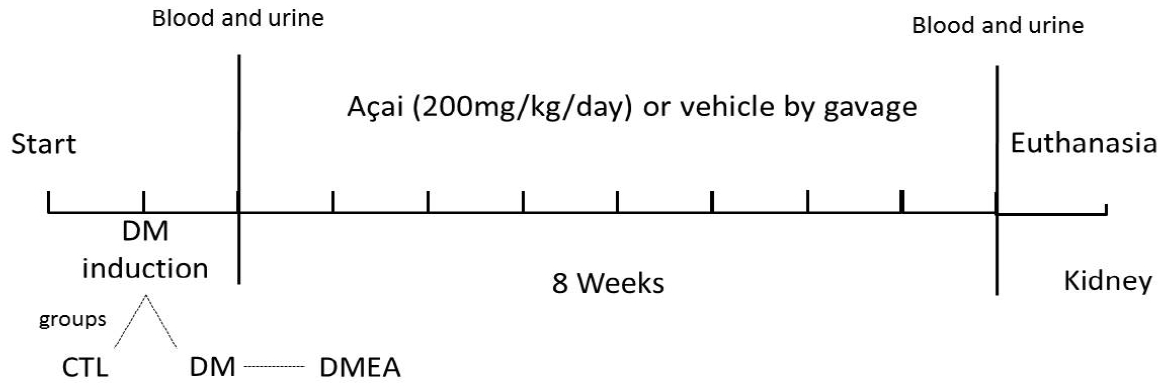

Flow diagram of the experimental protocol.

#### 2.2.5. Renal function

Plasma and urinary levels of creatinine were measured by colorimetric assay using a Labtest Creatinine kit (Centerlab Ltda, Sao Paulo, Brazil). The plasma urea concentrations were measured using a Labtest Urea CE kit (Centerlab Ltda, Sao Paulo, Brazil). The proteinuria was measured by colorimetric assay using a Sensiprot Labtest kit (Centerlab Ltda, Sao Paulo, Brazil) [37], and his analysis was performed in triplicate.

#### 2.2.6. Estimation of lipid peroxidation

Lipid peroxidation was estimated in plasma and urine at the end of the 8-week protocol using the TBARS method [38]. This analysis was performed in duplicate.

#### 2.2.7. Histological analysis

At the end of the 8-week protocol, half of each kidney was fixed in 10% formaldehyde, embedded in paraffin, sectioned to a 4 mm thickness and stained with periodic acid-Schiff reagent (PAS). The analysis was carried out at a magnification of 200× and analyzed by a pathologist, Dr Iria Visona, under blinded conditions.

#### 2.2.8. Statistical analysis

The results are expressed as the mean ± standard error of the media (SEM). First, the normality test (Kolmogorov-Smirnov) was performed, and then the data were analyzed. The differences among the groups were examined for statistical significance using one-way analysis of variance (ANOVA) followed by Newman–Keuls Multiple Comparison post-test for parametric data or Kruskal–Wallis followed by Dunn’s Multiple Comparison post-test for non-parametric data. For phenolic content analysis, linear

regression was used. Values were considered statistically significant when p < .05. Statistical analysis was performed in the program GraphPad Prism 5.0 (Graph Pad Software Inc., San Diego, USA).

## 3. Results

According to the analysis of the polyphenolic content, EA showed a very high concentration of total polyphenols and presented a particularly high capacity to reduce radical DPPH by 50%, compared to the literature (*Table 1*). Bonomo et al. [22] used the same brand of lyophilized product and showed that the extract contained 31.0 ± 2.4 mg/100 g of total monomeric anthocyanins, 8.8 ± 0.9 mg/100 g of cyanidin 3-O-glucoside and 8.7 ± 0.6 mg/100 g of cyanidin 3-O-rutinoside.

**Table 1.** Total phenolic and flavonoid contents and antioxidant activity of freeze-dried açai.

|  | Total phenolic content<br>(GAE mg/g) | Total flavonoid content<br>(QUE mg/g) | Antioxidant activity DPPH (IC<sub>50</sub><br>μg/mL<sup>-1</sup>) |
| --- | --- | --- | --- |
| Açai | 1344 ± 0.002 | 0.382 ± 0.232 | 22.79 |

3 The data are presented as the mean values ± SEM of triplicates (n = 3).

### 3.1. Assays in vitro

#### 3.1.1. Viability and cell proliferation

After treatment with EA, the viability evaluation of MiMC in NG medium showed that all groups had values above 90% of viable cells, showing no significant difference among the groups treated vs. NG, i.e., the non-cytotoxic effect of EA in MiMC.

There was a significant increase in cell proliferation in the HG group at all periods of treatment when compared to the NG group. The addition of 500 µg/mL EA significantly reduced the proliferation of MiMCs at all periods; 100 or 50 µg/mL EA also reduced the proliferation after 48 or 72 h compared to HG, p < .05 (**Fig. 1**).

**Fig. 1.**
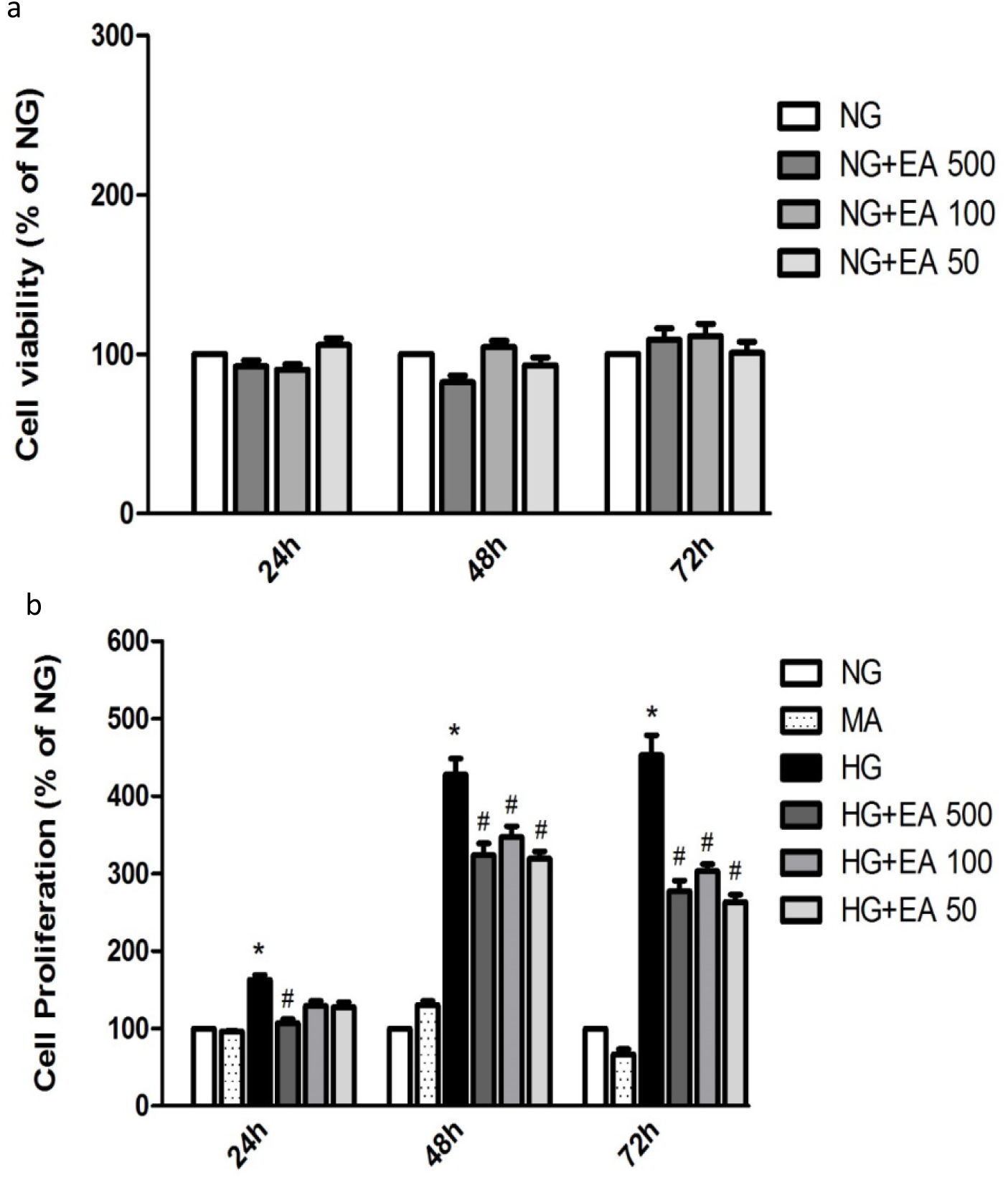
Cell viability (a) and proliferation (b) in MiMCs after 24, 48, or 72 h of EA treatment. NG: normal glucose; MA: mannitol; HG: high glucose; EA: açai extract. Mean ± SEM. One-way ANOVA with Newman-Keuls post-test, significance for p <.05: **\***vs. NG;^**#**^ vs. HG.

##### .Nitric oxide and ROS generation

NO (nmol/mL) determined by NOA in the cell supernatant (extracellular) was significantly increased in the HG group (66.8 ± 6.0) compared to the NG group (40.6 ± 2.4, p < .05) after 72 h of treatment. Supplementation with EA at doses of 500 µg/mL (40.8 ± 2.1) or 100 µg/mL (40.9 ± 1.8) resulted in a significant reduction in NO (p < .05), except for the HG + 50 group (45.2 ± 1.1), as shown in **Fig. 2a**.

**Fig. 2.**
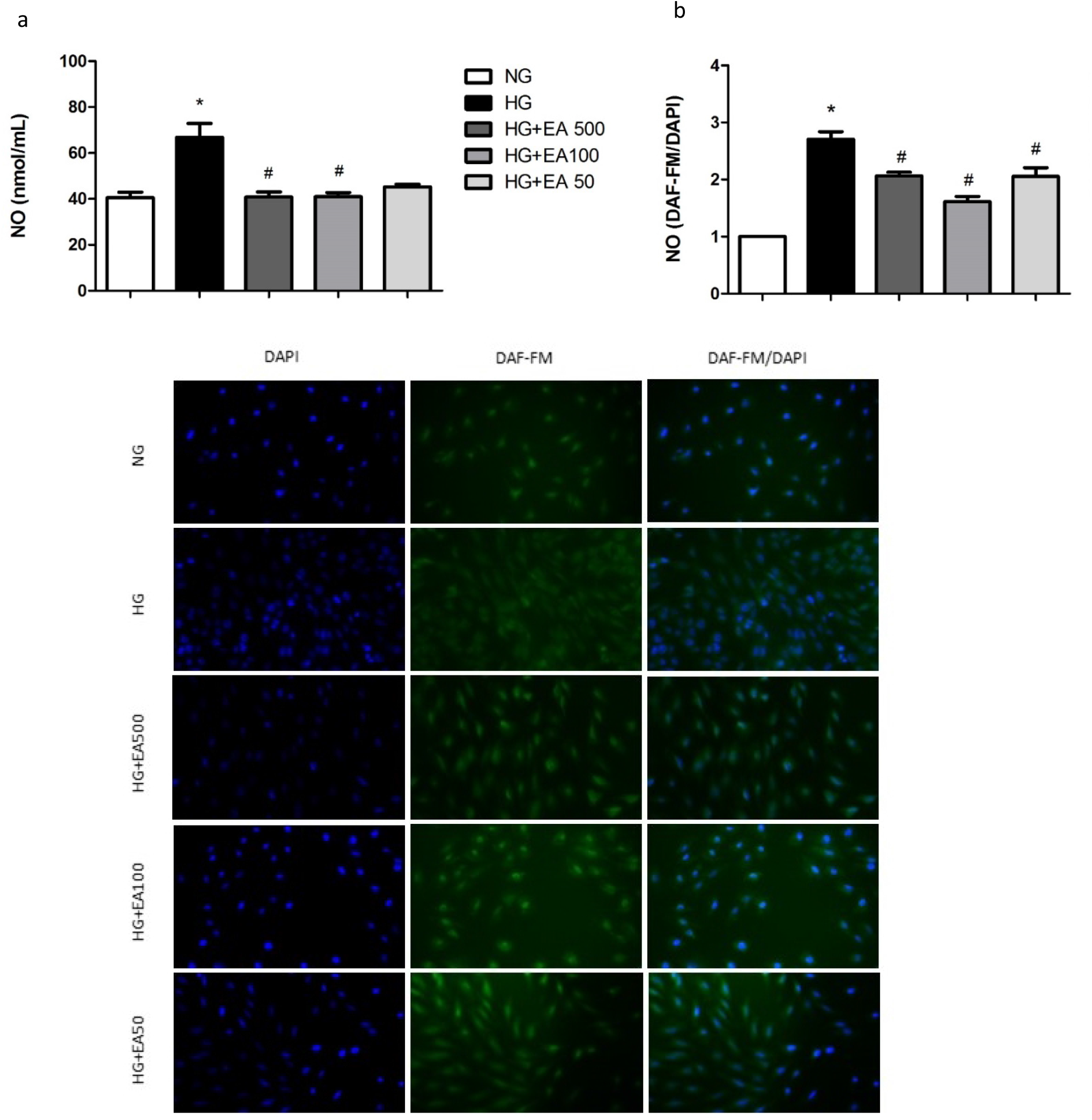
NO production (a) and NO (DAF-FM/DAPI) (b) staining after 72 h EA treatment in MiMCs. NO: nitric oxide. DAF-FM: diacetate 4-amino-5-methylamino-2’,7’- difluorofluorescein diacetate; DAPI: 4,6-diamidino-2-fenilindol. NG: normal glucose; HG: high glucose; EA: açai extract. Mean ± SEM. Kruskal-Wallis with Dunn’s post-test, significance for p < .05: **\***vs. NG;^**#**^vs. HG.

Similarly, the intracellular NO (DAF-FM/DAPI) fluorescence in MiMC showed that NO was increased in the HG vs. NG group (2.4 ± 0.1 vs. 1.0 ± 0.01 p < .05), indicating that these cells were sensitive to medium with high glucose stimulation. Treatment with EA at all doses significantly reduced the fluorescence (2.1 ± 0.1; 1.6 ± 0.1 and 1.9 ± 0.2, respectively, p < .05) in response to HG (**Fig. 2b**).

The ROS generation measured in the cells by the mean fluorescence intensity of DCF showed a significant increase in the HG group (6712 ± 771) compared to the NG group (3375 ± 333, p < .05) after 72 h. When the cells were treated with EA (500, 100 or 50 µg/mL), there was a significant reduction in the production of ROS (3925 ± 572; 3672 ± 482 and 3515 ± 562; respectively, p < .05) in relation to the HG group, as shown in **Fig. 3**.

**Fig. 3.**
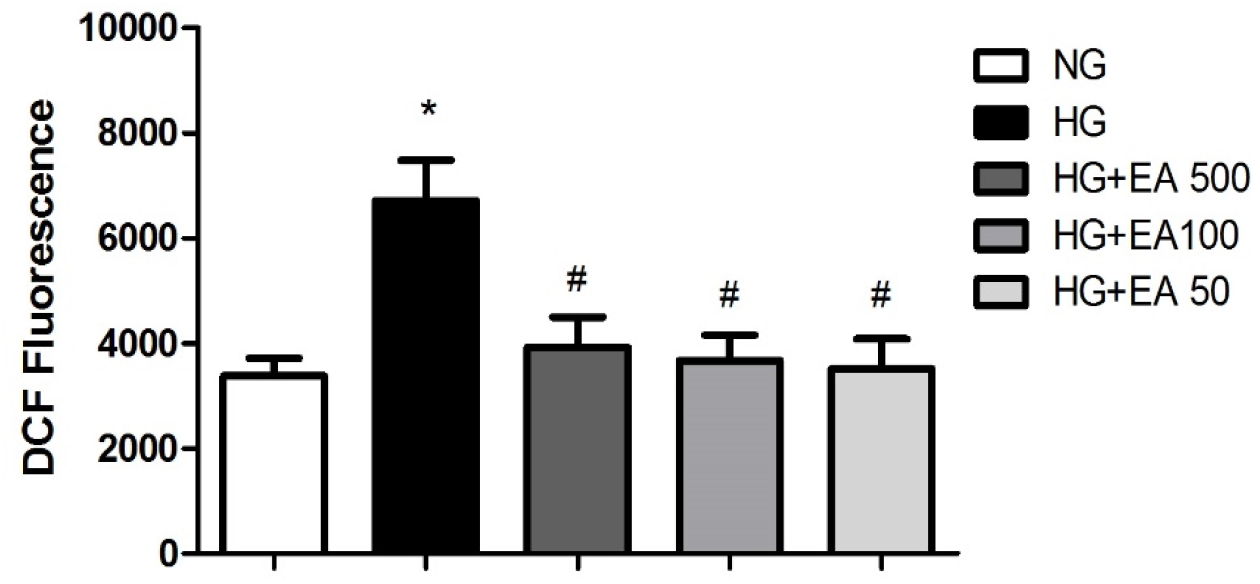
Intracellular ROS generation after 72 h EA treatment in MiMC. DCF: 2’,7’ dichlorodihydrofluorescein fluorescent. NG: normal glucose; HG: high glucose; EA: açai extract. Mean ± SEM. One-way ANOVA with Newman-Keuls post-test, significance for p < .05: **\***vs. NG;^**#**^vs. HG.

#### 3.1.3. EA effect on antioxidant expression

In **Fig. 4a**, there was a significant increase in EA 50 vs. HG (1.16 ± 0.1 vs. 0.83 ± 0.1, p < .05) in catalase.

**Fig. 4.**
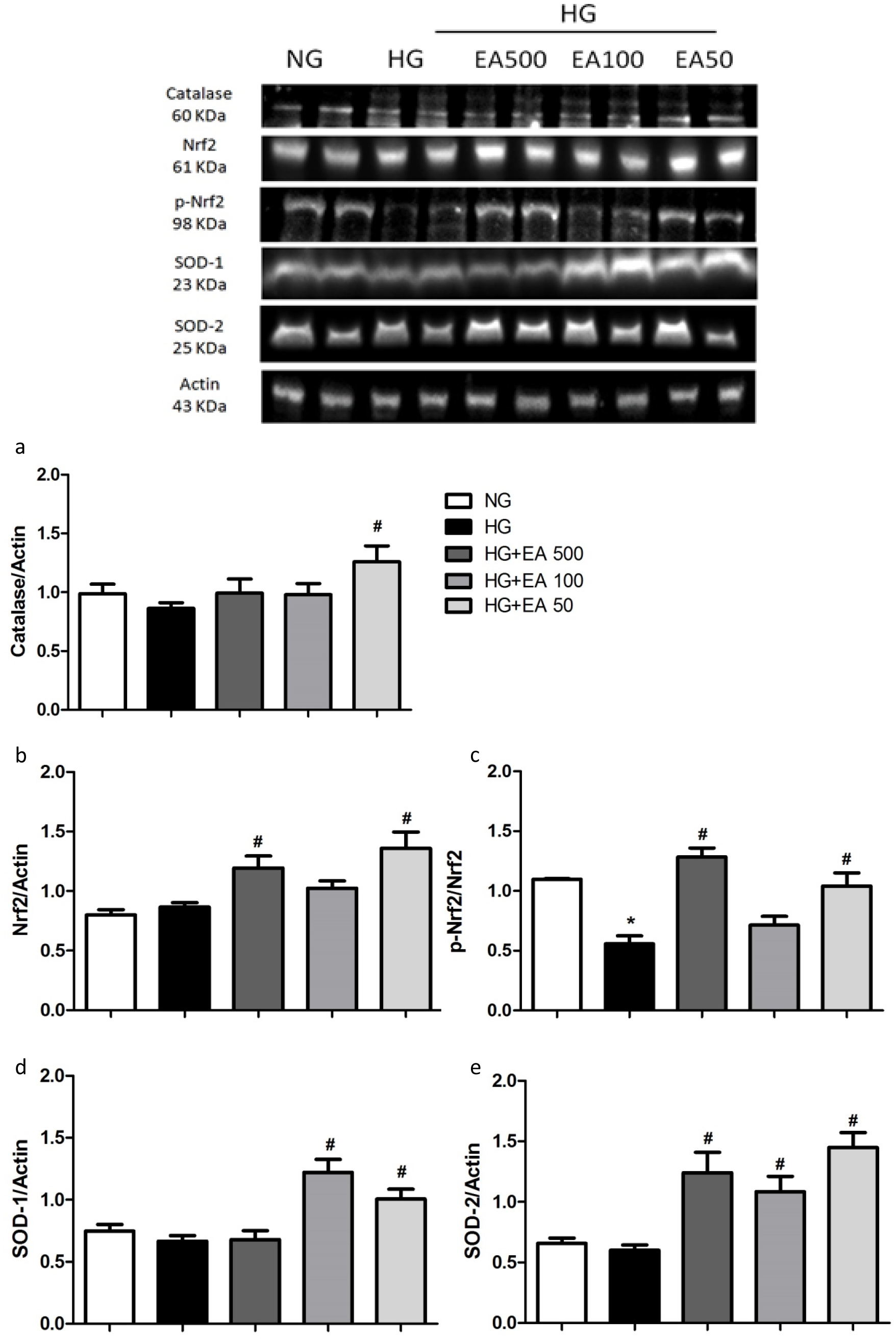
Quantification and representative image of catalase (a), Nrf2 (b), p-Nrf2/Nrf2 ratio (c), SOD-1 (d) and SOD-2 (e) after 72 h EA treatment in MiMC. NG: normal glucose; HG: high glucose; EA: açai extract; Nrf2: nuclear factor erythroid-derived 2; p-Nrf2: phosphorylated; SOD1: superoxide dismutase 1 – cytosolic; SOD-2: superoxide dismutase 2-mitochondrial. Mean ± SEM. One-way ANOVA with Newman-Keuls post-test or Kruskal-Wallis with Dunn’s post-test, significance for p < .05: **\***vs. NG;^**#**^vs. HG.

Nrf2 expression increased significantly in the EA 500 and EA 50 groups vs. HG (1.17 ± 0.1 and 1.26 ± 0.1 vs. 0.79 ± 0.1, p < .05). Phosphorylated Nrf2 (p-Nrf2) was significantly decreased in the HG vs. NG (0.56 ± 0.1 vs. 1.1 ± 0.0, p < .05); however, EA 500 and EA 50 (1.3 ± 0.1 and 1.0 ± 0.1, respectively, p < .05) had a significant increase in their expression vs. HG (**Fig. 4b,c**).

SOD-1 showed a significant increase in EA 100 and 50 µg/mL vs. HG (1.22 ± 0.1 and 1.09 ± 0.1 vs. 0.67 ± 0.04, p < .05), while SOD-2 increased significantly in all EA doses vs. HG (1.22 ± 0.1; 1.07 ± 0.1 and 1.48 ± 0.1, respectively, p < .05), as seen in **Fig. 4d, e**.

#### 3.1.4. EA effect on inflammatory profile

iNOS was significantly increased in HG vs. NG (0.99 ± 0.1 vs. 0.67 ± 0.04, p < .05); in contrast, EA significantly reduced the protein content of this isoform in the HG + EA 100 or 50 groups (0.58 ± 0.1 and 0.59 ± 0.1, p < .05) in relation to HG, as shown in **Fig. 5a**.

**Fig. 5.**
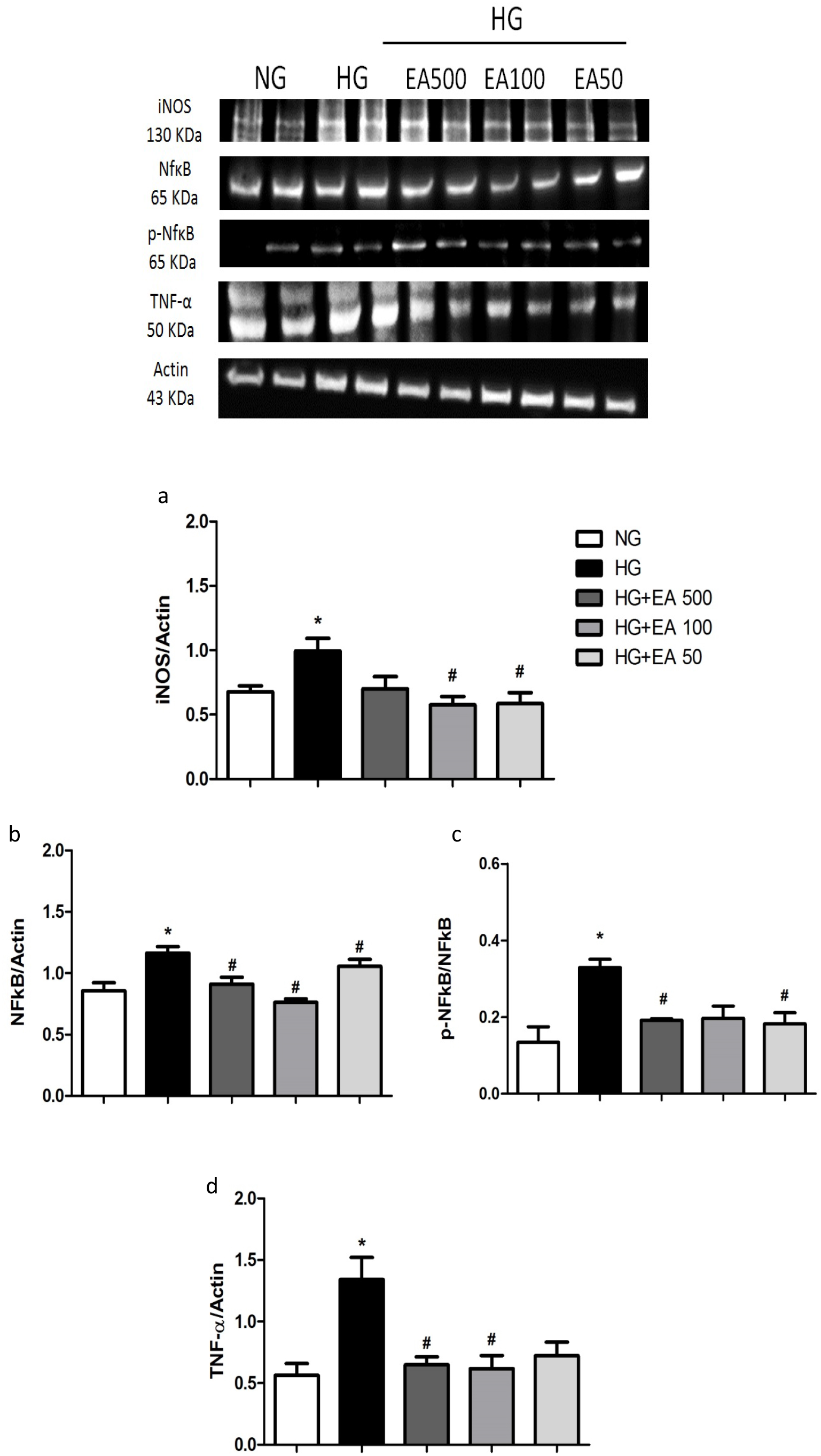
Quantification and representative image of iNOS (a), NF-κB (b), p-NF-κB/NF-κB ratio (c) and TNF-α (d) after 72 h EA treatment in MiMC. iNOS: inducible nitric oxide synthase; NF-κB: nuclear transcription factor; p-NF-κB: phosphorylated NF-κB; NG: normal glucose; HG: high glucose; EA: açai extract. Mean ± SEM. One-way ANOVA with Newman-Keuls post-test or Kruskal-Wallis with Dunn’s post-test, significance for p < .05: **\***vs. NG;^**#**^vs. HG.

NF-κB p65 expression was greater in HG vs. NG (1.16 ± 0.1 vs. 0.83 ± 0.1, p < .05); however, there was a significant reduction after treatment with all EA doses (0.91 ± 0.1; 0.8 ± 0.04 and 1.03 ± 0.1, respectively, p < .05) compared to the HG group (**Fig. 5b**).

In its phosphorylated form, p-NF-κB expression was greater in HG in relation to the control (0.33 ± 0.02 vs. 0.13 ± 0.04, p < .05); the higher and lower (0.19 ± 0.0 and 0.18 ± 0.0, p < .05) doses, EA 500 and EA 50, respectively, were reduced compared to the HG group (**Fig. 5c**).

TNF-α expression in the HG group presented a significant increase compared to NG (1.34 ± 0.2 vs. 0.56 ± 0.1, p < .05); however, after 72 h of treatment with EA, there was a significant decrease of approximately 50% in both the EA 500 and 100 µg/mL groups (0.65 ± 0.1 and 0.61 ± 0.1, respectively, p < .05) vs. HG, as seen in **Fig. 5d**.

### 3.2. Assays in vivo

#### 3.2.1. Metabolic data, renal function and lipid peroxidation in animals after EA treatment

According to the protocol, the rats presented the classic symptoms of diabetes when compared to control animals; the group that received EA showed significant improvement in body weight recovery vs. DM, p < .05. The diabetic vs. control group revealed significant differences in plasma urea, creatinine, and proteinuria; on the other hand, the DMEA vs. DM group showed a significant reduction in these parameters, demonstrating an improvement in renal function (*Table 2*).

**Table 2.** Metabolic profile, analysis of renal function and lipid peroxidation at the 8th week of treatment.

| Parameters | CTL | DM | DMEA |
| --- | --- | --- | --- |
| Fasting Glucose (mg/dL) | 99.7 ± 1.4 | 557.9 ± 9.3 * | 506.7 ± 5.6 |
| Liquid Intake (mL/24 h) | 30.0 ± 0.6 | 143.6 ± 7.4 * | 130.0 ± 5.0 |
| Chow Intake (g/24 h) | 20.5 ± 0.7 | 32.6 ± 0.9 * | 32.7 ± 1.1 |
| Diuresis (mL/24 h) | 12.5 ± 0.9 | 98.2 ± 3.9 * | 98.5 ± 2.0 |
| Body Weight (g) | 382.7 ± 4.5 | 191.8 ± 10.4* | 248.3 ± 17.3 # |
| Plasma Creatinine (mg/dL) | 0.51 ± 0.01 | 0.96 ± 0.07 * | 0.73 ± 0.05 # |
| Plasma Urea (mg/dL) | 16.46 ± 0.34 | 22.64 ± 0.98 * | 17.11 ± 0.43 # |
| Proteinuria (nmol/24 h) | 2.57 ± 0.23 | 12.68 ± 1.21 * | 5.74 ± 0.56 # |
| TBARS Excretion (nmol/24 h) | 71.9 ± 6.3 | 301.1 ± 11.5 * | 267.9 ± 9.6 # |
| Plasma TBARS (nmol/mL) | 2.82 ± 0.05 | 3.19 ± 0.25 | 2.96 ± 0.12 |

CTL: control group; DM: diabetic group; DMEA: diabetic treated with açai; TBARS: thiobarbituric acid reactive substances. The results are presented as the mean ± SEM. *One-way* ANOVA with Newman-Keuls post-test or Kruskal-Wallis with Dunn post-test. N= 4-5 per group. p < .05: \*vs. CTL; # vs. DM.

As shown in Table 2, TBARS excretion was significantly increased in the DM vs. CTL group; EA significantly reduced lipid peroxidation in the DMEA group compared to the DM group. Plasma TBARS levels were not significantly different between groups.

##### Histological analysis in the renal cortex

The histological analysis by light microscopy and staining with periodic acid-Schiff reagent staining (PAS) revealed some changes, such as diffuse sclerosis and enlargement of the mesangial matrix in the glomeruli (black arrows) and tubules with glycosidic degeneration (black circles) in the diabetic groups; however, there was a notable reduction in these changes in DMEA vs. DM, as shown in **Fig. 6**.

**Fig. 6.**
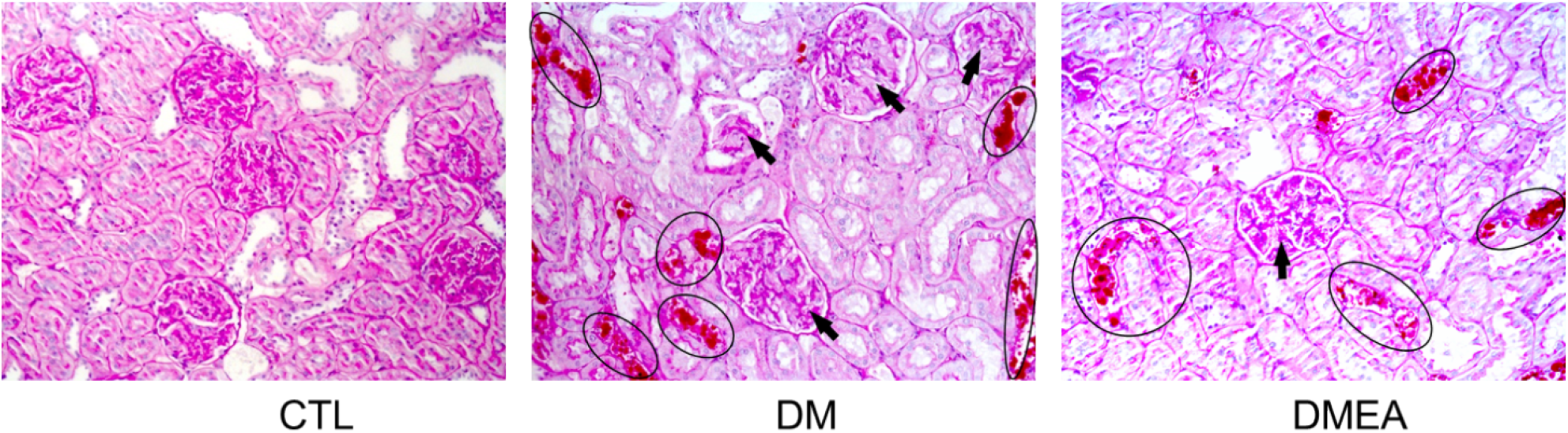
Microscopic analysis of renal tissue sections after the 8th week of EA in animals. CTL: control group; DM: diabetic group and DMEA: diabetic treated with açai. Glomeruli with diffuse sclerosis and increased mesangial matrix (black arrows) and tubules with glycosidic and hyaline degenerations (black circles).

## 4. Discussion

In the present study, we showed that açai extract (EA), rich in polyphenols, reduced cell proliferation, NO levels and ROS production in MiMCs treated with high glucose. In contrast, EA in these cells was able to increase the expression of proteins from the antioxidant defense system and reduce inflammatory factors. To our knowledge, this is the first study to use açai as a nonpharmacological way to control the deleterious effects of high glucose in mesangial cells.

Açai has been studied due to its high content of anthocyanins and flavonoids; some studies have shown beneficial effects of açai in some experimental models of diseases. *In vitro* and *in vivo* studies showed that açai has antioxidant, anti-inflammatory, antiproliferative and antitumorigenic properties, as well as pro-apoptotic and cholesterol lowering action [24, 39, 40]. The benefits of açai on health are already partially proven, and this antioxidant effect was also found in other fruits and vegetables [41]; Pacheco-Palencia et al. [42] showed that polyphenolic extracts of various fruits and vegetables have an inhibitory effect on cell proliferation.

Açai, in addition to being known for its antioxidant potential, has approximately 2.4% protein and 5.9% lipids; monounsaturated oleic acid is the primary fat (56.2%), palmitic acid (24.1%) and linoleic acid (12.5%) [20, 43]. Anthocyanins are generally considered the main contributors to the antioxidant activity of the pulp; the pulp has 48.6 μmol Trolox equivalents/ml of antioxidant capacity. In addition, other phenolic compounds are present in the fruit, such as 3-glycosides of anthocyanin, ferulic acid, epicatechin, p-hydroxybenzoic acid, gallic acid, protocatechuic acid, catechin, ellagic acid, vanillic acid, and p-coumaric acid. and gallotannins [43], which confers its high polyphenolic content.

Similar to other fruits, such as cranberries, blueberries, and strawberries, açai has a high antioxidant capacity. A study evaluated the total polyphenol content (w/w% GAE) and antioxidant activity (DPPH, IC50; g/mL) of crude berry fruit extracts rich in polyphenols and anthocyanins, such as blackberry (6.8% /1968.6), black raspberry (4.1% /2865.9), blueberry (9.8% /381.1), cranberry (7.7% /392.6), red raspberry (6.4%/ 268.5), and strawberry with 3.8% of total polyphenols [44].

Another study with carcinogenic cells treated with açai hydroalcoholic extract in three doses (10, 20, or 40 μg/mL) for 24, 48 or 72 h did not show cytotoxicity in 24 or 48 h [45], corroborating our data, once we did not observe cytotoxicity after treatment with EA, in MiMC at these periods of time.

High glucose induces cell proliferation and collagen synthesis through mechanisms involving the increase in ROS and transforming growth factor beta 1 (TGF-β1) in mesangial cells [46]. Liu et al [47] found increased proliferation of rat mesangial cells in HG medium compared to NG. In our study, exposure to high glucose also promoted cell proliferation, independent of osmolarity (as shown in the mannitol group), and increased ROS generation; these effects were suppressed by EA, suggesting that ROS are important mediators of high glucose-induced mesangial cell activation.

Song et al [48] treated mesangial cells with high glucose medium and delphinidin (an anthocyanin) and obtained similar results, in which anthocyanin reduced cell proliferation, ROS generation, suppressed NOX-1 mRNA, and mitochondrial superoxide. Hogan et al [39] also showed an inhibitory effect of açai extract (50, 100 or 200 μg/mL) on the proliferation of C6 rat brain glioma cells, showing that the higher the dose, the greater the reduction of cell proliferation. The açai extract was recently reported as a dose-dependent inhibitor of proliferation in HT-29 colon carcinoma cells. The polyphenol fractions of açai pulp have shown a reduction in leukemic HL-60 cell proliferation by activating caspase-3, inducing cell apoptosis [49].

Our results demonstrated that high glucose generated oxidative stress through increased ROS production and extracellular and intracellular measured NO overproduction in MiMC, probably via NF-κB and iNOS stimuli. El Remessy et al [50] suggested that NO generation can be stimulated by an increase in glucose levels. Hyperglycemia, via glucose oxidation, can stimulate the production of ROS, including the superoxide anion, which reacts with NO to form peroxynitrite, a highly cytotoxic nitrogen reactive species. By increasing intracellular oxidative stress, AGEs activate the transcription factor NF-κB, promoting the upregulation of various target genes controlled by this factor and increasing NO production, which is believed to be a mediator of β-cell damage [50]. Other researchers showed that exposure of mouse mesangial cells to HG (25 mM) stimulated iNOS gene and protein expression and enhanced NO synthesis [51]; similar data were shown in our study, demonstrating in turn that EA reduced the inflammatory stimuli in 72 h.

The activation of the NF-κB pathway may trigger pro-or anti-apoptotic cascades, predominantly pro-apoptotic cascades [52]. Inhibition of this pathway protects pancreatic β-cells from cytokine-induced apoptosis *in vitro* and *in vivo* and can be a potential strategy for protecting these cells [53]. High glucose generates ROS in mesangial cells and upregulates NF-κB, as shown by the elevation of the p-NF-κB/NF-κB ratio. NF-κB-dependent pathways play an important role in the infiltration of macrophages and kidney damage [54]. In addition, a study showed that NF-κB is also regulated by angiotensin II; this regulates cell proliferation, apoptosis, fibrosis and the inflammatory response through NF-κB-dependent pathways [55]. Furthermore, açai pulp extracts rich in anthocyanins showed a reduction in oxidative stress and inflammation via inhibiting iNOS, cyclooxygenase-2 (COX-2), p38 mitogen-activated protein kinase (p38-MAPK), TNFα and NF-κB, attenuating the release of extracellular NO [56].

TNF-α, an inflammatory cytokine, activates the proinflammatory signaling pathway of NF-κB [57]. There are two pathways for NF-κB activation: the classic pathway (regulated by activation IkB kinase β - IKKβ), which is triggered by toll-like receptors (TLRs) and proinflammatory cytokines such as TNF-α and interleukin-1, leading to activation of RelA [58], and the alternative pathway that is activated by lymphotoxin β, resulting in activation of the RelB/p52 complex but not TNF-α. Another study showed that supplementation with anthocyanins inhibited NF-κB activation induced by lipopolysaccharides (LPS) in monocyte culture and further reduced TNF-α in healthy volunteers [59]. Our results showed a significant increase in the expression of NO, iNOS, TNF-α and the p-NF-κB/NF-κB ratio in the HG group, characterizing the activation of inflammatory mediators; however, all these markers were reduced after treatment with EA, possibly via NF-κB inactivation.

In our study, EA increased the expression of catalase, both cytosolic Cu-Zn-SOD (SOD-1) and mitochondrial Mn-SOD (SOD-2), which are regulated by Nrf2 activation. Under normal conditions, Nrf2 is linked with Keap1 (the protein in the cytoplasm), but under oxidative and inflammatory conditions, Nrf2 decouples from Keap1 and translocates to the nucleus, where it activates the expression of several cytoprotective proteins and enzyme genes, increasing cellular survival. Therefore, Keap1-Nrf2-ARE signaling plays an important role in maintaining the cellular redox balance [60].

Açai modulated ROS production by neutrophils in the liver of rats with diabetes and increased the antioxidant defense system [61]. In another study, açai reduced the levels of malondialdehyde and carbonyl protein in hypertensive rats, recovered the levels of SOD, CAT and GPx and the expression of SOD-1 and SOD-2, preventing vascular remodeling; these findings suggest that açai produced an antihypertensive effect and prevented endothelial dysfunction and vascular structural alterations [62]. Additionally, an anthocyanin extract study showed an increase in antioxidant defense system enzymes (SOD, CAT and GPx) providing protection against H2O2 and glucose toxicity in β-pancreatic cells [63], corroborating our study.

Cyanidin-3-glucoside, a component present in açai, reduced light-induced retinal oxidative stress by activating the Nrf2 antioxidant pathway and attenuated the expression of inflammation-related genes by suppressing NF-κB activation in an animal experimental model [64]. In another study, rats fed a açai-enriched diet presented Nrf2 modulation and ubiquitin-proteasomal pathway regulation in the hippocampus and frontal cortex, resulting in benefits to cognitive function during brain aging [17]. Our data agree with these studies, since EA treatment promoted an upregulation of Nrf2, resulting in a significant increase in all antioxidants, suggesting that açai was able to modulate the Nrf2 antioxidant response in conditions of increased oxidative stress and inflammation via NF-κB activation.

Consistent with previous reports [21, 65], we found that oxidative stress was elevated in DN. In the present study, the oxidative damage assessed by TBARS was markedly increased in the DM group, and NO and ROS production was elevated in the HG group. EA significantly reduced these oxidative stress markers, indicating an important antioxidant effect that could contribute to renal protection.

Due to hyperglycemia, the reabsorption of excess filtrate leads to renal overload, triggering podocyte detachment, death of epithelial and mesangial cells and intense production of the ECM, which can cause glomerular sclerosis and contribute to the progression of DN [2]. In our study, alterations in parameters of renal function, such as elevated plasma creatinine and urea, in association with proteinuria characterized DN; however, the decline in renal function was prevented by EA, which was observed through the reduction of histological markings in the DMEA group.

Açai extract protected the kidneys of spontaneously hypertensive rats with STZ-induced diabetes, reducing albuminuria excretion, serum levels of urea and creatinine and renal fibrosis, evidenced by decreased TGF-β and collagen IV [66]. In another study, açai preserved kidney morphology and function in hypertensive rats with chronic ischemic renal injury (2K1C), preventing albuminuria and increasing serum urea and creatinine levels, contributing to the reduction of glomerular damage [67]; these findings agree with our research, since we observed a significant improvement in renal function in diabetic animals that consumed EA. As a limitation of this study, we think that the creatinine and urea levels in urine would need to be determined in these animals.

Our findings allow us to conclude that EA was beneficial to delay diabetic kidney disease, as observed by the preservation of both renal function and morphology, suggesting that the mechanisms underlying these effects of EA may involve the reduction of cell proliferation, oxidative stress, ROS, NO, lipid peroxidation and inflammatory mediators via NF-κB inactivation and upregulation of the Nrf2 antioxidant response pathway.

## 5. Conclusion

In the present study, açai extract was able to decrease the proliferation and oxidative stress induced by high glucose in MiMC by suppressing inflammatory mediator signaling (NF-κB - iNOS - TNF-α) and, at the same time, upregulating the Nrf2 antioxidant response pathway, conferring a delay in the progression of renal disease and resulting in protection of the mesangial cells. Despite not significantly reducing fasting glycemia, EA helped in the recovery of body weight, reduced oxidative stress and improved renal function, demonstrating less glycosidic degeneration in the renal tissue of these animals compared to DM.

Further studies are needed to evaluate the açai mechanisms of action in the progression of diabetes, which could be useful to control this disease.

## Acknowledgements

The authors acknowledge all the members of our laboratory for their technical assistance. This study was supported by Conselho Nacional de Desenvolvimento Cientifico e Tecnologico (CNPq) and Fundacao de Amparo a Pesquisa do Estado de Sao Paulo, #2014/26750-9 (FAPESP).

